# Detection of oat crown rust disease in Taiwan (2019-2021)

**DOI:** 10.1101/2024.03.12.584714

**Authors:** Chung-Ying Ho, Eva C Henningsen, Ssu-Tung Chen, Hiran A. Ariyawansa, Eric S. Nazareno, Jana Sperschneider, Peter N. Dodds, Jakob Riddle, Shahryar F. Kianian, Melania Figueroa, Yung-Fen Huang

**Author notes:** Corresponding author: Y-F Huang.

## Abstract

Oat is a minor forage crop grown in Taiwan. Only a few historical records of oat rust disease have been reported in the country, therefore the pathogen population remains poorly characterized. A rust-like disease outbreak was detected at the Experimental Farm of National Taiwan University in 2019, which caused significant damage to the field experiments. To determine the identity of the pathogen responsible for this disease outbreak, we collected infected foliar material. Disease signs suggested infection by the oat crown rust fungus. Hence, common procedures in rust pathology were applied to confirm the identity of the pathogen with phenotypic and molecular diagnostic techniques. A total of 50 field samples from infected oat cultivars were collected in 2019 and five rust isolates were purified in 2020 and 2021. Phylogenetic analysis based on ITS sequences indicated that the pathogen was likely *Puccinia coronata* f. sp. *avenae* (*Pca*), which was further supported by the placement of Taiwanese isolate NTU-01 with other *Pca* representatives in a phylogenetic tree of Basidiomycete fungi. Phenotyping assays across 36 oat differential lines demonstrated that Taiwanese isolates are phenotypically similar with relatively limited virulence. This study presents the first molecular confirmation of *Pca* in Taiwan and reports the virulence profiles of Taiwanese *Pca* population.

## Introduction

Oat (*Avena sativa* L.) ranks as the seventh largest cereal in the world in terms of grain production and cultivated surface (FAOSTAT, 2022). Oat has desirable health benefits in both human food and as forage for rumen (Butt et al. 2008; Coblentz et al. 2013; Contreras-Govea and Albrecht 2006). The most significant foliar disease of oat worldwide is crown rust (Nazareno et al. 2018). Oat crown rust disease is caused by the Basidiomycete biotrophic fungus *Puccinia coronata* f. sp. *avenae* (*Pca*), which can cause grain yield losses between 10 to 40% and total loss in severe epidemics (Nazareno et al. 2018). *Pca* also reduces the forage dry matter yield and quality (Andrzejewska et al. 2019). The life cycle of *Pca* is heterocious, with asexual reproduction occurring on oat and the sexual cycle occurring on several species of *Rhamnus* (Nazareno et al. 2018). In places where no alternative host is reported, it is often assumed that clonality (asexual reproduction) and somatic mutation drive the evolution of pathogen and the emergence of new strains (Figueroa et al. 2020)

The use of resistant varieties is the most economically effective and sustainable approach to control rust diseases (McCallum et al. 2007; Periyannan et al. 2017). Like other rust systems, resistance to *Pca* can be classified into race-specific or race-nonspecific categories (Ohm and Shaner 1992; Periyannan et al. 2017). Race-specific resistance, also known as seedling resistance, follows the classical gene-for-gene interaction where immunoreceptors encoded by the resistance genes in the plant (*R*) recognize the avirulence factors (*Avr*) in the pathogen (Dodds 2023; Flor 1970). For oat crown rust, these *R* genes are referred to as *Pc* genes. Race-specific resistance has been largely deployed in rust resistance breeding programs around the globe. However, the lifespan of released oat varieties that carry single *R* genes (*Pc* genes) does not often last longer than five years because of the rapid virulence evolution of *Pca* populations frequent emergence of new virulent strains of the pathogen (Carson 2011; Nazareno et al. 2018). Assignment of races of *Pca* follows scoring of infection types in a set of differential lines where each line is postulated to carry different *Pc* resistance genes (Nazareno et al. 2018). Unfortunately, the oat crown differential set has not been genetically characterized at depth. Across different oat crown rust differential sets, phenotypic and genotypic discrepancies among common lines have been identified, suggesting that race assignments are not always comparable across institutions (Nguyen et al. 2023). Furthermore, other studies highlight the potential redundancy of crown rust resistance genes within oat lines (Hewitt et al.; Miller et al. 2020).

Non-specific race resistance, also known as adult plant resistance (APR) or partial resistance, does not necessarily manifest at the seedling stage and often presents delayed symptom development or low disease severity at adult plant stage (Nazareno et al. 2018). This resistance is more durable but individual loci have smaller effects and varying mechanisms for conferring resistance. Efforts to characterize adult plant resistance and new sources of race-specific resistance and combine them for oat breeding programs are ongoing (Admassu-Yimer et al. 2018; Babiker et al. 2015; Díaz-Lago et al. 2003; Leonard 2002; Nazareno et al. 2022; Nazareno et al. 2018).

In Taiwan, oat is a minor crop that is grown for cut-and-carried forage by dairy cattle farmers (Chen and Huang 2021). Although Taiwan is far from the center of origin of oat and oat seed is mostly imported from other countries, wild oat (*Avena fatua* L.) is present in some places across Taiwan (iNaturalistTW 2024). The tropical/subtropical environment of Taiwan requires that spring oat varieties typically grown in temperate regions are sown between October to December and harvested between February and April to avoid heat stress. Since the 1980’s, oat was grown annually in small parcels at the Experimental Farm of National Taiwan University (NTU) to support the oat breeding program. Larger forage oat field experiments were planted in NTU annually beginning in 2015.

In the growing season of 2017 – 2018, we observed rust-like symptoms in the field for the first time, which were followed by a disease outbreak in the season 2018 – 2019. While corn and sugarcane rusts have been reported in Taiwan throughout the broader rust research community (Hooker 1985; Hou et al. 1978; Hsieh et al. 1977; Purdy 1985), records of oat rusts did not attract much attention as they were written in Japanese or in Chinese and reported only observation sites and pathogen names (Chen et al. 1980; Sawada 1928, 1943). Given the poor characterization of cereal rusts in Taiwan and the recently emerging need to introduce rust resistance traits into our oat breeding program, we isolated the pathogen from diseased field trials from 2019-2021 and phenotyped 55 rust samples on a set of oat crown rust differential lines. We then verified the identity of the pathogen with phylogenetic trees generated from ITS and whole-genome sequencing data. Here, we confirm through infection assays and phylogenetic analysis that *Pca* is the causal agent of the 2019 disease epidemic detected in oat fields on the Experimental Farm of NTU.

## Materials and Methods

### Puccinia coronata f. sp. avenae collection, storage, and microscopy

*Pca* samples were collected in 2019, 2020, and 2021 from the Experimental Farm, College of Bio-resources and Agriculture, NTU (25.01514, 121.54008). In April 2019, 50 samples of rust-infected leaves were collected from 43 oat lines (Supplemental Table 1). Infected leaves were dried at room temperature in an airtight container with desiccant for one week. The 50 dried leaf samples were shipped to the USDA-ARS Cereal Disease Laboratory (CDL), Saint Paul, MN, USA for oat differential line inoculation and virulence testing. In May and June 2020 and February 2021, samples of *Pca* urediniospores were directly collected from infected leaves using gelatin capsules, among which five isolates were purified using standard single-pustule technique (Supplemental Table 1). Urediniospores were dried at room temperature in an airtight container with desiccant for one week. Capsules containing dried urediniospores were stored at −80°C before use. A heat-shock activation at 45°C for 10 minutes (Miller et al. 2020) was applied to the −80°C-stored urediniospores prior to the inoculation. For urediniospore morphology observation, fresh urediniospores were sampled from infected leaves and mixed with distilled water for brightfield observation using Olympus System Microscope BX51 (Olympus, Tokyo, Japan).

### Plant materials and growth conditions

A subset of 36 lines from the oat differential set were used for the virulence testingboth at the USDA-ARS CDL and NTU (Chong et al. 2000; Nazareno et al. 2018). In addition to the oat differential set, two highly susceptible oat cultivars in Taiwan, Swan and NTU Sel. No.1, were used for the purification and amplification of rust isolates for experiments conducted at NTU.

Oat seeds were disinfected with 1% NaClO for five minutes, rinsed by deionized water three times, and then soaked in deionized water at room temperature for 16 hours before planting. Six seeds were sowed in the medium of peat moss (pH 5.5, Kekkilä-BVB, Vantaa, Finland) and red soil mixture (1:1) in a 3-inch round pot for *Pca* purification and amplification while seeds were sowed in line in 3-inch square pots for pathogenicity evaluation. Oat seedlings for purification and amplification were treated with 3 mM maleic hydrazide (Sigma-Aldrich, St. Louis, MO, USA) at emergence. Approximately seven to nine days after germination, seedlings reached one- to two-leaf stage were used for inoculation.

### Plant inoculation and pathogenicity evaluation

Pathogenicity evaluation with the 2019 samples was completed at the USDA ARS CDL facility as described by Miller et al. (2021). Bulk inoculation was used for each sample without undergoing single pustule purification. Infection type was scored with the 0-4 scale adapted from Murphy (1935) where scores from 0 to 2 indicate resistance and scores from 3 to 4 indicates susceptibility. Pathogen isolation and characterization undertaken in Taiwan followed a similar methodology to characterize isolates collected in 2020 and 2021. The plant inoculation and pathogenicity method used in Taiwan was modified as follows. Fresh urediniospores generated from a single pustule culture were suspended in Isopar M (ExxonMobil, Houston, TX, USA) at 1-mg spores per ml (Cabral and Park 2014). To ensure an even inoculation on the leaves, we used a sprayer (K-3A-05, diameter 0.5 mm, Kinki Factory, Osaka, Japan) to spread the inoculum at a pressure of 12.5 pound per square inch. The sprayer was set 30 cm away in parallel to the leaves. After inoculation, the mineral oil was allowed to evaporate for 40 minutes, then the inoculated plants were placed in an opaque plastic box which provided a dark and humid environment (90% relative humidity) at 18-20°C for 16 hours. After 16 hours, the inoculated plants were placed in a greenhouse at 20/16°C. Ten to twelve days after inoculation, infection type was scored based on five leaves per line using a modified Cobb scale from 0 to 4 (Supplemental Table 2) (Cabral and Park 2014; Miller et al. 2020; Murphy 1935; Nazareno et al. 2018). Infection types were further converted into 0 and 1 to indicate avirulent (infection types from 0 to 2) and virulent (infection types of 3 and 4) reactions for heatmap visualization using the *pheatmap()* function implemented in R 4.3.1 (R-Core-Team 2023). Manhattan distance was used to cluster similar isolates.

### DNA extraction from rust isolates, ITS amplification and phylogenetic analysis

The CTAB method (Doyle and Doyle 1990) was used to extract DNA from rust urediniospores for the amplification of the internal transcribed spacer (ITS). We use forward primer ITS-1F (5′-CTTGGTCATTTAGAGGAAGTAA-3′) and reverse primer ITS4 (5′-TCCTCCGCTTATTGATATGC-3′) to amplify ITS (Gardes and Bruns 1993; Manter and Vivanco 2007; White et al. 1990). Polymerase Chain Reaction (PCR) was performed in a solution containing 12.5 μl 2×Taq Master Mix (Protech, Taipei, Taiwan), 1 μl of each primer at 10 μM, with 1 μl DNA and 9.5 μl ddH2O added up to 25 μl with the following program: initial heating of 95°C for 5 minutes, 32 cycles of 95°C for 30 seconds, 55°C for 30 seconds, 72°C for 60 seconds, and a final extension at 72°C for 10 minutes. The PCR products were separated by electrophoresis with 2% agarose gel under 110V for 30 minutes. PCR product was purified using GenepHlow™ Gel/PCR Kit (DFH300, Geneaid, New Taipei City, Taiwan). The purified PCR products were sequenced by Sanger sequencing (ABI3730, Core Laboratory of Biotechnology, National Taiwan University). The sequencing data were curated using CodonCode Aligner (CodonCode Corporation, Centerville, Ohio, USA) prior to phylogenetic analysis. Seventeen *Puccinia coronata* ITS sequences and two *Puccinia graminis* ITS sequences (Szabo 2006) in addition to one ITS from *Chrysomyxa conituberculata* (Wang et al. 2022) were downloaded from NCBI and were included in the phylogenetic analysis. Phylogenetic analysis was performed using the maximum likelihood method implemented in software Mega X (Kumar et al. 2018) with 5,000 bootstrap. *C. conituberculata* was used as the outgroup.

### Whole-genome sequencing and phylogenetic analysis of Taiwanese Pca isolate NTU1

For whole-genome sequencing, genomic DNA was extracted using the Omniprep DNA isolation kit (G-Biosciences, St. Louis, MO, USA) from 20 mg of urediniospores of *Pca* isolate NTU1. DNA concentration was determined using Equalbit 1 × dsDNA HS Assay Kit (Vazyme, Nanjing, China) and a Qubit 3.0 Fluorometer (Life Technologies, Singapore) before submission for whole-genome sequencing. DNA sequencing was completed at Azenta Life Sciences (Suzhou, China) with NovaSeq S4, 300 cycles (Illumina, San Diego, CA, USA) to produce 150 bp paired-end reads with 35X target coverage. Read2tree was used to place NTU1 within *Puccinia* (Dylus et al. 2024). Briefly, 1149 marker genes for 52 Basidiomycete species representing 39 genera were downloaded from the Orthologous Matrix (OMA) browser and prepared for use in read2tree (read2tree -- reference) (Altenhoff et al. 2020). Illumina reads from 11 isolates representing seven *Puccinia* species/formae specialis (*P. graminis* f. sp. *avenae*, *P. graminis* f. sp. *tritici*, *P. polysora*, *P. coronata* f. sp. *avenae*, *P. striiformis* f. sp. *tritici*, *P. hordei*, *P. triticina*) and *Melampsora larici-populina* were aligned to the downloaded Basidiomycota marker genes (read2tree --reads) (Supplemental Table 3). Subsequently, the alignments were merged and a phylogenetic tree was constructed (read2tree --merge_all_mappings --tree). Bootstrap values (n = 1000) were added with IQ-TREE v2.2.0.5 (iqtree -m LG -B 1000) (Minh et al. 2020).The resulting tree was visualized with iTOL (Letunic and Bork 2021).

## Results

### Collection of P. coronata f. sp. avenae in Taiwan

The signs and symptoms present in oat trials at the Experimental Farm of NTU from 2019 to 2021 and spore morphology suggested that the pathogen was likely *P. coronata* f. sp. *avenae* (*Pca*), the oat crown rust fungus (Fig. 1). Macroscopic signs included yellow-orange sporulation covering primarily leaves (Figure 1A and 1B). This observation aligns with signs of oat crown rust rather than oat stem rust, which is darker orange in color and forms larger oblong pustules on leaves and stems (Martens 1985; Simons 1985). Spores collected from infected plants are circular and approximately 25 µM in diameter as reported previously for *Pca* urediniospores (Savile 1984).

**Figure 1.**
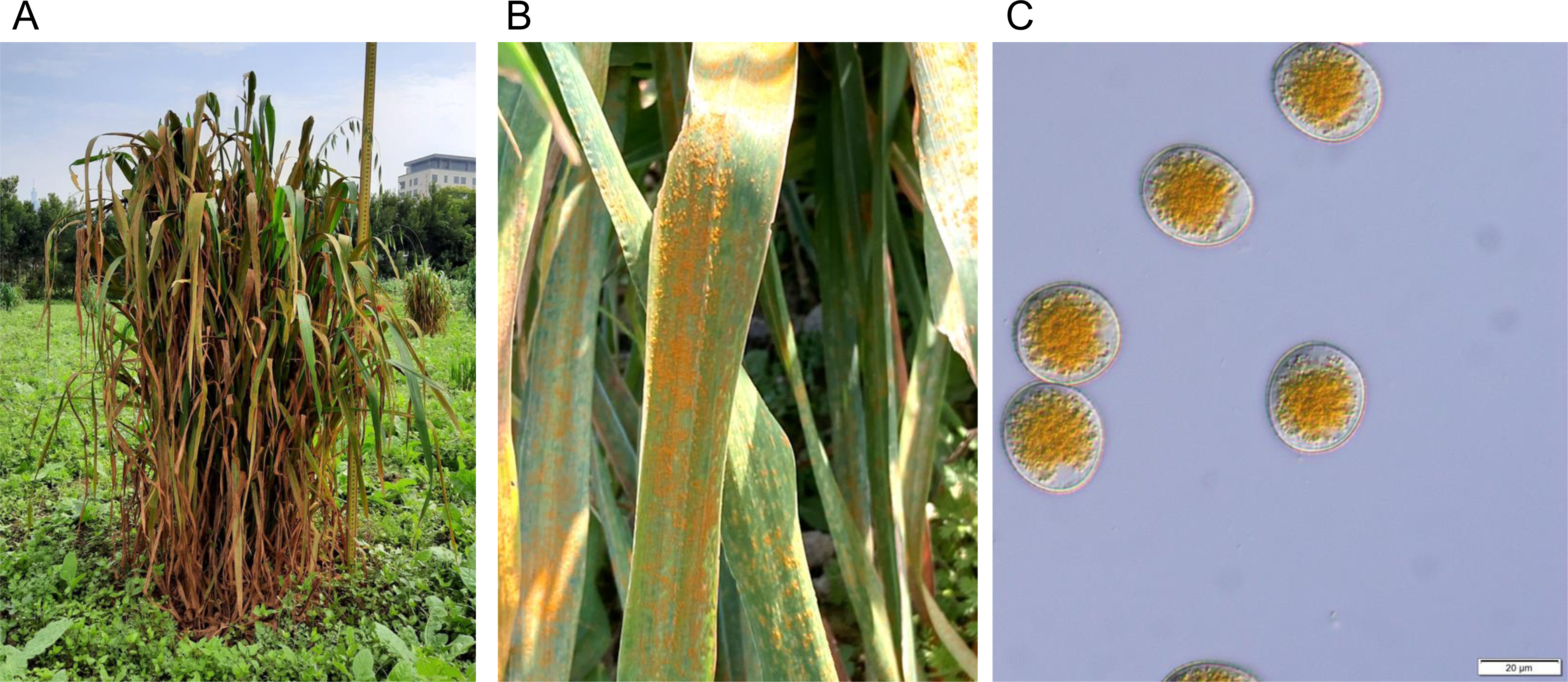
Photos of infected plant materials from 2021 at the Experimental Farm, National Taiwan University. (A) infected plants in the field. (B) a closer view of sporulating pustules on oat leaves. (C) urediniospores collected from infected tissue examined with microscopy at 400X magnification.

We revived 50 samples from 2019 and 5 samples from 2020-2021 and tested them across 36 oat differential lines routinely used for race assignment at the USDA-ARS CDL (Nazareno et al. 2018). The phenotypes of Taiwanese *Pca* samples collected from 2019-2021 were similar, with six sets of isolates having identical phenotypes across the 36 differential lines (Figure 2). Only 17 of the 36 differential lines used showed variation in virulence across isolates, and 11 of these lines differed for six or fewer isolates (Pc14, Pc38, Pc39, Pc48, Pc52, Pc54, Pc59, Pc60, Pc61, IAB605Xsel., TAM-O-405) (Figure 2). Most isolates were virulent to Pc36, Pc56, Pc67, Pc96, and Marvelous. Virulence was not detected for 19 lines (Pc40, Pc45, Pc46, Pc50, Pc51, Pc55, Pc57, Pc58, Pc62, Pc63, Pc64, Pc68, Pc91, Pc94, WIX4361-9, Belle, HiFi, Leggett, Stainless).

**Figure 2.**
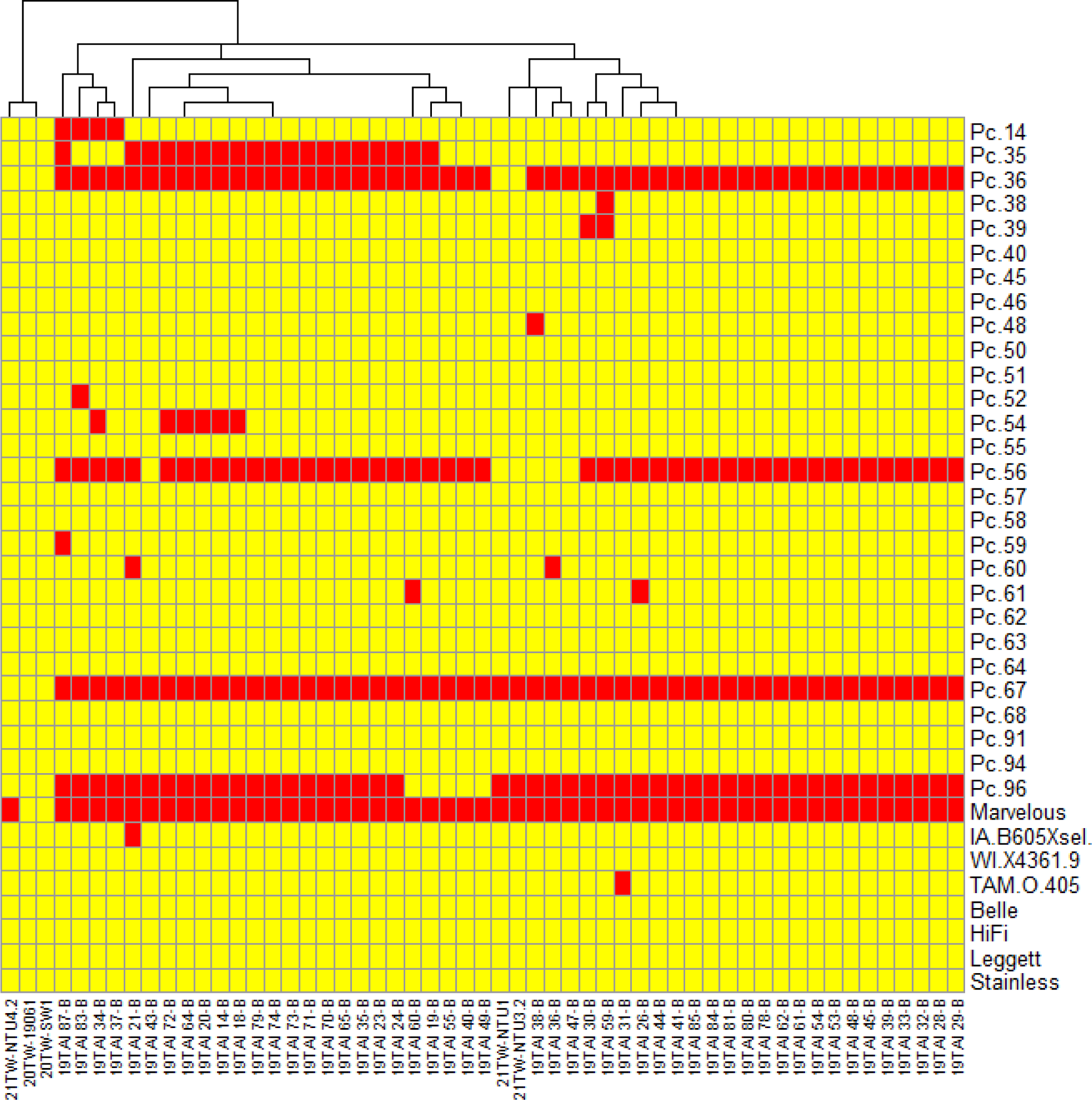
Heatmap of virulence phenotypes from 55 *Puccinia coronata* f. sp. *avenae* isolates collected in Taiwan on 36 oat differential lines. Collection year is indicated by the first two numbers of isolate names. Colors indicate virulence (red) or avirulence (yellow). Isolates are clustered by phenotypic similarity.

Marvelous is used as a control line for phenotyping in the North America as it consistently displays an extremely susceptible phenotype against crown rust. However, in the infection assays performed in Taiwan for 2020-2021 isolates, the phenotype of Marvelous was borderline, considered avirulent for some isolates and virulent for others. Lines Swan and NTU Sel. No. 1 had a more severe susceptible phenotype when tested in Taiwan, like what was observed for 2019 isolates tested on Marvelous in the USA. Therefore, Swan may be better than Marvelous for use as a susceptible check in tropical and subtropical climates.

### Molecular confirmation of the identity of P. coronata f. sp. avenae

ITS sequences amplified from four Taiwanese rust isolates and published sequences from *Puccinia graminis* and *P. coronata* f. sp. *avenae* were used to generate a phylogenetic tree, with the ITS from *Chrysomyxa contibuerculata* used as a Pucciniomycete outgroup (Szabo 2006; Wang et al. 2022). The four ITS sequences from Taiwanese *Pca* isolates were nearly identical to *P. coronata* ITS collected on oat, *Lolium*, and *Rhamnus cathartica*, which correspond to previously described Clade V (representing *Pca*) in *P. coronata* subspecies trees (Liu and Hambleton 2013; Szabo 2006). As further confirmation, whole genome sequencing data for Taiwanese isolate NTU1 was used to place the isolate in a Basidiomycete phylogenetic tree. This phylogeny accurately delineated the major Basidiomycota subdivisions Agaricomycotina, Wallemia, Ustilagomycotina, and Pucciniomycotina with 100% bootstrap support (Figure 4). NTU1 was placed with *Pca* isolates 12SD80, 12NC29, and Pca203 with high bootstrap support (100%) (Figure 4). This phylogeny does not place NTU1 with *P. graminis* f. sp. *avenae*, another significant oat rust pathogen. The available genetic evidence suggests *Pca* caused the 2019 oat rust epidemic in Taiwan, showcasing the first robust identification of *Pca* in Taiwan.

**Figure 3.**
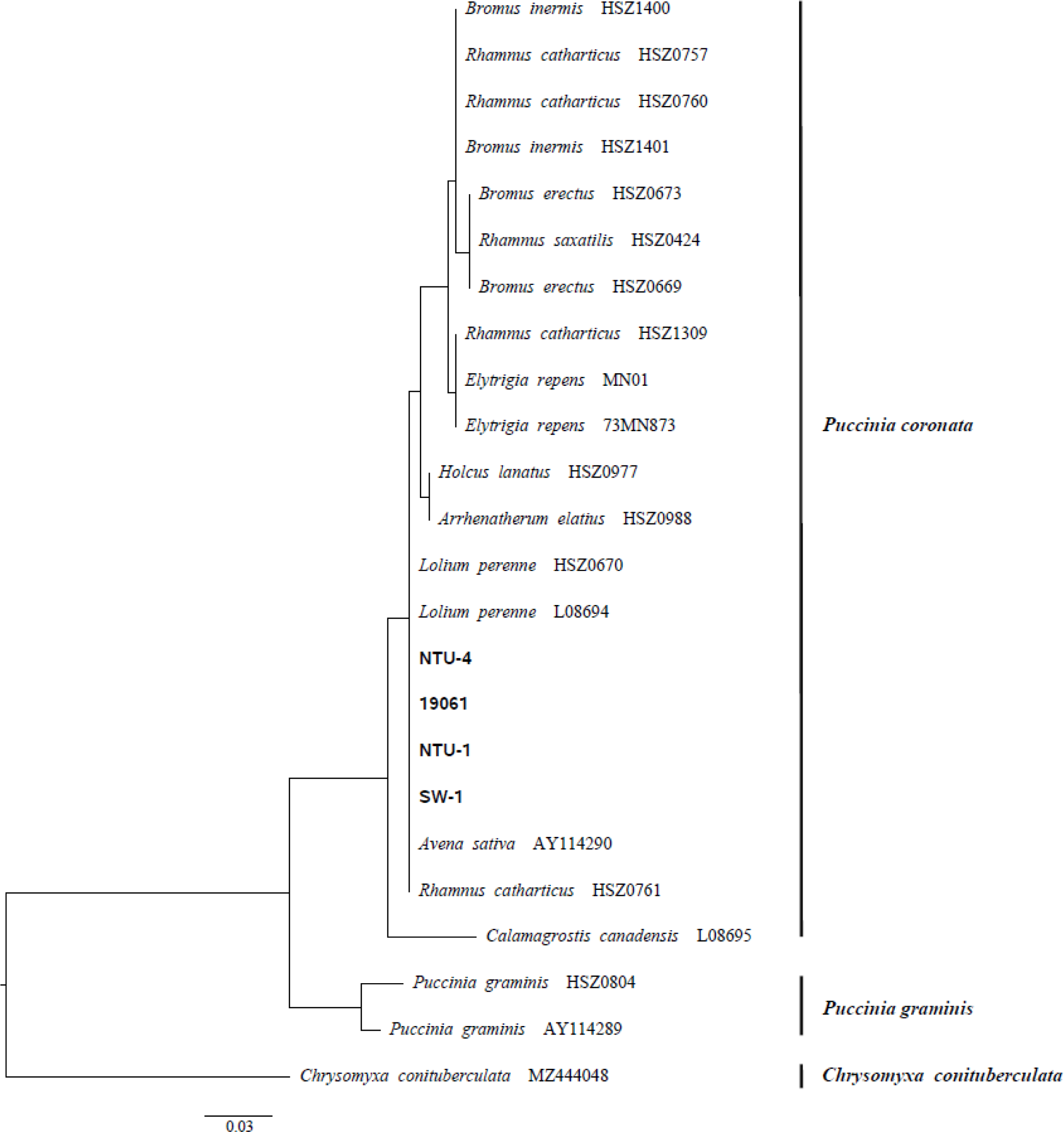
Midpoint rooted phylogenetic tree based on published *Puccinia coronata* f. sp. *avenae* (*Pca*) ITS sequences, including *P. graminis* f. sp. *tritici* for comparison and *Chrysomyxa conituberculata* as an outgroup. Taiwanese *Pca* isolates are highlighted with bold text. Tree scale is mean substitutions per site.

**Figure 4.**
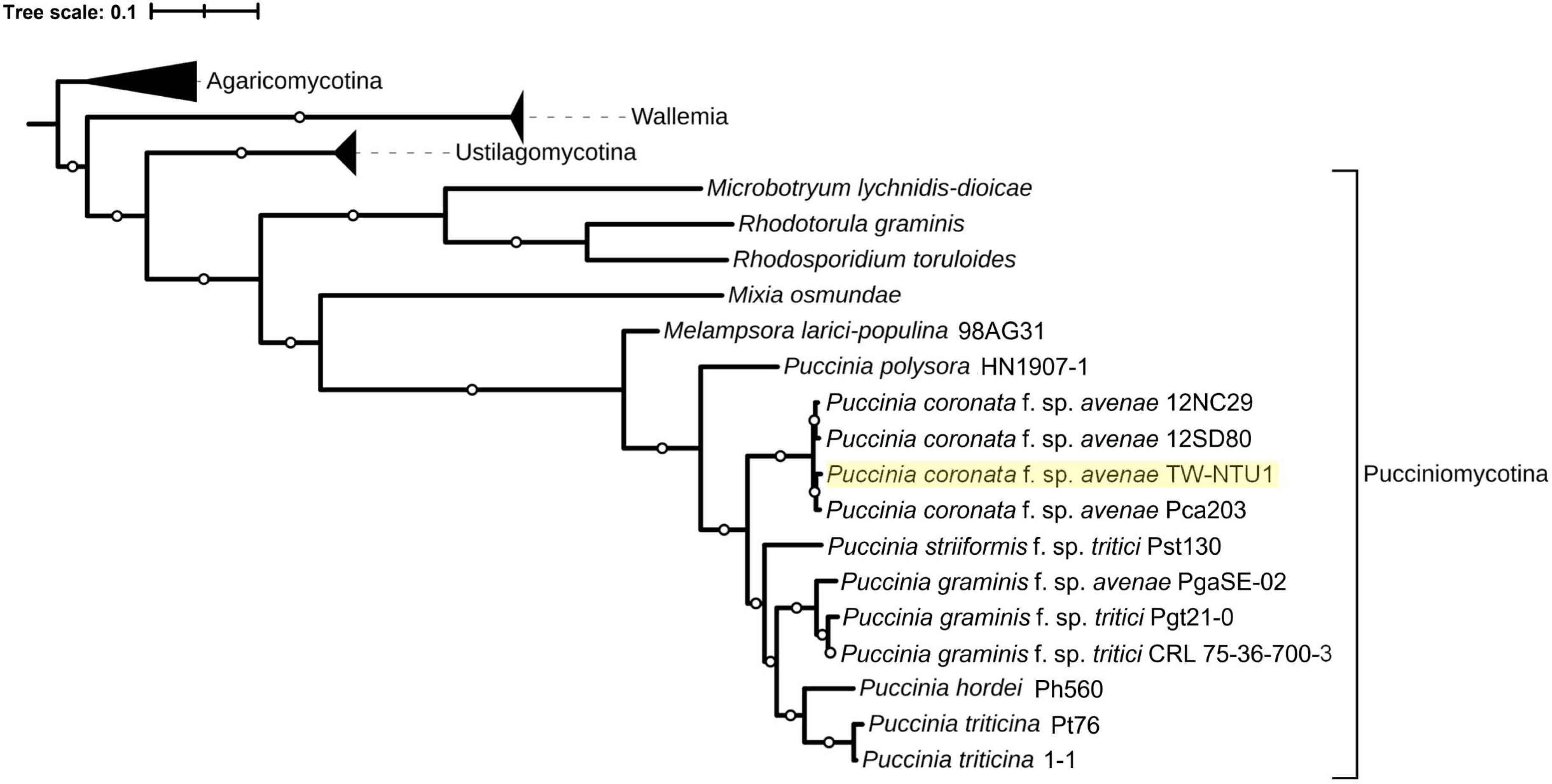
Midpoint rooted phylogenetic tree of Basidiomycete species from the Orthologous Matrix browser and publicly available rust short read data generated from alignments between 1149 marker genes. The rust isolate from Taiwan is highlighted in yellow. Basidiomycota subdivisions other than Pucciniomycotina (shown with bracket) are collapsed. 1000 bootstraps were performed and branches with 100% support have white circles at their midpoint. Tree scale is mean substitutions per site.

## Discussion

Rust fungi described as oat pathogens (*Avena* spp.) include oat crown rust (*Pca*) and oat stem rust (*Pga*). Although oat is not known to host other *P. coronata formae specialis*, other grass genera such as *Hordeum* (barley), *Lolium*, and *Phalaris* have been reported infected with *Pca* (Liu and Hambleton 2013). Further, the best-described alternate host for *Pca* (*Rhamnus cathartica*) and other *Rhamnus* species host multiple *P. coronata formae specialis*. Because of these overlapping host ranges and complicated subspecies relationships, it is prudent to confirm the identity of rust growing on oat. The macroscopic signs and phylogenetic trees generated from ITS and marker gene sequences confirm that *Pca* as the cause of recent rust epidemics in Taiwan over *Pga* or other *Puccinia* spp. Although rust on oats has been described in Taiwan in the past, previous reports lack the required information to validate the presence of *Pca* specifically (Chen et al. 1980; Sawada 1928, 1943), positioning this publication as the first report of *Pca* with molecular evidence in Taiwan.

While the cause of a sudden *Pca* epidemic in Taiwan is not certain, it is possible that warmer conditions across south and eastern Asia reported during the winter 2018/2019 and monsoon cycle facilitated appropriate climatic conditions (e.g., temperature, precipitation) for *Pca* epidemic development (Sato 2019).

The oat cropping area in Taiwan declined from 440 hectares in 1982 to 9.64 hectares in 2019 (Agriculture 2024), suggesting that the epidemic is not likely to be related to large-scale changes in oat cropping area or practices. However, the authors note that from 2015 oat cultivation area increased on the NTU experimental farm. It is possible that this local increase in susceptible oats enabled the epidemic growth of a small existing *Pca* population in the area. Regardless of what caused the first significant outbreak of *Pca* in Taiwan, oat crown rust was also sampled in 2020 and 2021 and has been observed in subsequent years. *Pca* will likely persist in Taiwan unless conditions that prevent uredinial reproduction (host availability, weather) occur over an extended period to cause local extinction.

Compared to *Pca* populations from the USA and Australia, Taiwanese samples collected in 2019 have relatively limited virulence traits (Henningsen et al. 2023; Hewitt et al. 2023; Miller et al. 2020). In the USA, virulence to all resistance sources has been detected (Hewitt et al. 2023), whereas virulence was either absent or rarely detected in the 2019 Taiwan population for all but eight lines. Many of these tested resistance sources have been overcome in Australia at moderate to high frequency as well, with a virulence to a few differential lines being rare (Pc58, Pc59, Pc94, HiFi, Leggett, Stainless, WIX4361-9, Belle, TAM-O-405) or absent (Pc63) like in Taiwan (Henningsen et al. 2023). Given the phenotypic similarity and low virulence of Taiwanese isolates, it is likely that the *Pca* isolates responsible for the epidemic in Taiwan are members of an older, more avirulent race. However, this hypothesis should be examined with whole-genome sequencing of additional Taiwanese isolates, placement in a species-specific phylogenetic tree, and generation of a haplotype phased genome reference.

There were some notable differences in phenotyping conducted in the USA (2019 population) as compared to in Taiwan (2020-2021 isolates). For example, virulence for Pc36 and Marvelous were fixed in the 2019 population, however 2020-2021 isolates were all avirulent on Pc36 and two of the five were avirulent on Marvelous. These results may accurately reflect the variation between isolates collected across these years. Alternatively, this could arise from systemic biases that complicate comparing phenotyping results across countries and institutions. Individual researchers make biased assessments during visual phenotyping. Germplasm purity is another challenge, as recent work has described genotypic and phenotypic inconsistencies between the same oat differential lines from different seed sources (Nguyen et al. 2023). Further, oat lines with multiple sources of resistance may begin segregating following their initial release, further complicating the interpretation of phenotyping conducted at different institutions (Nguyen et al. 2023). Finally, resistance loci effective against other rust fungi display temperature sensitivity (Adhikari et al. 2000; Gousseau et al. 1985; Niu et al. 2014; Yu et al. 2023). These, among other, factors, may partially explain the differences between 2019 assays conducted in the USA and 2020-2021 assays conducted in Taiwan.

## Acknowledgments

The authors are grateful to the technical assistance of the Core Laboratory of Biotechnology, National Taiwan University, for ITS Sanger sequencing and the Azenta NGS Laboratory in Suzhou for NGS sequencing. This study was funded by National Science and Technology Council of Taiwan (Grant number: 109-2313-B-002-028-MY3) and internal CSIRO funding. ECH was supported by the ANU University Research Scholarship and ANU/CSIRO Digital Agriculture PhD Supplementary Scholarship.

