## Supplemental Tables for "Detection of oat crown rust disease in Taiwan (2019-2021)"

Supplemental Table 1

| Sample | Oat line | Collected by | Date collected | Location | Taiwan_field_ID |
|---|---|---|---|---|---|
| 19TAI 14-B | PI436085_375 | Yung-Fen Huang | 2019-04-25 | NTU experimental farm | B7C2R9 |
| 19TAI 18-B | PI264853_788 | Yung-Fen Huang | 2019-04-25 | NTU experimental farm | B7C1R11 |
| 19TAI 19-B | PI264853_788 | Yung-Fen Huang | 2019-04-25 | NTU experimental farm | B7C1R11 |
| 19TAI 20-B | PI264853_788 | Yung-Fen Huang | 2019-04-25 | NTU experimental farm | B7C1R11 |
| 19TAI 21-B | PI573536WIR_1926 | Yung-Fen Huang | 2019-04-25 | NTU experimental farm | B7C3R11 |
| 19TAI 23-B | Kanota | Yung-Fen Huang | 2019-04-25 | NTU experimental farm | B7C1R10 |
| 19TAI 24-B | PI577928_71_4 | Yung-Fen Huang | 2019-04-25 | NTU experimental farm | B6C7R11 |
| 19TAI 26-B | PI287442AVE_136_59 | Yung-Fen Huang | 2019-04-25 | NTU experimental farm | B7C4R9 |
| 19TAI 28-B | PI344849_54 | Yung-Fen Huang | 2019-04-25 | NTU experimental farm | B7C4R12 |
| 19TAI 29-B | PI344849_54 | Yung-Fen Huang | 2019-04-25 | NTU experimental farm | B7C4R12 |
| 19TAI 30-B | HA05AB42-20 | Yung-Fen Huang | 2019-04-25 | NTU experimental farm | B7C6R9 |
| 19TAI 31-B | Ajay | Yung-Fen Huang | 2019-04-25 | NTU experimental farm | B8C1R9 |
| 19TAI 32-B | Ajay | Yung-Fen Huang | 2019-04-25 | NTU experimental farm | B8C1R9 |
| 19TAI 33-B | CILLA | Yung-Fen Huang | 2019-04-25 | NTU experimental farm | B8C1R11 |
| 19TAI 34-B | CILLA | Yung-Fen Huang | 2019-04-25 | NTU experimental farm | B8C1R11 |
| 19TAI 35-B | Lamont | Yung-Fen Huang | 2019-04-25 | NTU experimental farm | B8C1R10 |
| 19TAI 36-B | PI183990_122 | Yung-Fen Huang | 2019-04-25 | NTU experimental farm | B8C2R10 |
| 19TAI 37-B | 00Ab7006 | Yung-Fen Huang | 2019-04-25 | NTU experimental farm | B8C2R8 |
| 19TAI 38-B | PI158139CI_4817 | Yung-Fen Huang | 2019-04-25 | NTU experimental farm | B8C3R9 |

| | | | | | |
|---|---|---|---|---|---|
| 19TAI 39-B | PI158139CI_4817 | Yung-Fen Huang | 2019-04-25 | NTU experimental farm | B8C3R9 |
| 19TAI 40-B | PI573546WIR_2137 | Yung-Fen Huang | 2019-04-25 | NTU experimental farm | B8C4R9 |
| 19TAI 41-B | PI662093TJK2006_086 | Yung-Fen Huang | 2019-04-25 | NTU experimental farm | B8C4R10 |
| 19TAI 43-B | PI374407_55_71_67 | Yung-Fen Huang | 2019-04-25 | NTU experimental farm | B8C6R9 |
| 19TAI 44-B | PI159130_42_3 | Yung-Fen Huang | 2019-04-25 | NTU experimental farm | B8C5R13 |
| 19TAI 45-B | PI177861Bayrak_Sumbuleli | Yung-Fen Huang | 2019-04-25 | NTU experimental farm | B8C3R13 |
| 19TAI 47-B | Nudist | Yung-Fen Huang | 2019-04-25 | NTU experimental farm | B8C3R12 |
| 19TAI 48-B | PI168115_4100 | Yung-Fen Huang | 2019-04-25 | NTU experimental farm | B6C4R7 |
| 19TAI 49-B | PI168115_4100 | Yung-Fen Huang | 2019-04-25 | NTU experimental farm | B6C4R7 |
| 19TAI 53-B | Swan | Yung-Fen Huang | 2019-04-26 | NTU experimental farm | B7C7R5 |
| 19TAI 54-B | PI168068_1610 | Yung-Fen Huang | 2019-04-26 | NTU experimental farm | B8C1R3 |
| 19TAI 55-B | PI60753_1250 | Yung-Fen Huang | 2019-04-30 | NTU experimental farm | B1C3R7 |
| 19TAI 59-B | PI286414 | Yung-Fen Huang | 2019-04-30 | NTU experimental farm | B2C3R9 |
| 19TAI 60-B | PI405739VI_30 | Yung-Fen Huang | 2019-04-30 | NTU experimental farm | B2C4R12 |
| 19TAI 61-B | HA08-03X45-1 | Yung-Fen Huang | 2019-04-30 | NTU experimental farm | B3C1R10 |
| 19TAI 62-B | PI573548WIR_4953 | Yung-Fen Huang | 2019-04-30 | NTU experimental farm | B3C1R11 |
| 19TAI 64-B | PI287340AVE_148_59 | Yung-Fen Huang | 2019-04-30 | NTU experimental farm | B3C1R13 |
| 19TAI 65-B | Optimum | Yung-Fen Huang | 2019-04-30 | NTU experimental farm | B3C1R14 |
| 19TAI 70-B | PI573567WIR_10950 | Yung-Fen Huang | 2019-04-30 | NTU experimental farm | B4C3R1 |
| 19TAI 71-B | PI405735VI_26 | Yung-Fen Huang | 2019-04-30 | NTU experimental farm | B4C5R10 |
| 19TAI 72-B | PI577966_135b_5 | Yung-Fen Huang | 2019-04-30 | NTU experimental farm | B4C5R12 |
| 19TAI 73-B | Brooks | Yung-Fen Huang | 2019-04-30 | NTU experimental farm | B4C5R13 |

| | | | | | |
|---|---|---|---|---|---|
| 19TAI 74-B | PI168121_4315 | Yung-Fen Huang | 2019-04-30 | NTU experimental farm | B4C6R8 |
| 19TAI 78-B | PI266280WIR_6103 | Yung-Fen Huang | 2019-04-30 | NTU experimental farm | B5C2R11 |
| 19TAI 79-B | Sang | Yung-Fen Huang | 2019-04-30 | NTU experimental farm | B5C6R3 |
| 19TAI 80-B | PI344833_39 | Yung-Fen Huang | 2019-04-30 | NTU experimental farm | B5C7R10 |
| 19TAI 81-B | PI577876_29a_1_2 | Yung-Fen Huang | 2019-04-30 | NTU experimental farm | B5C3R8 |
| 19TAI 83-B | PI266267WIR_5396 | Yung-Fen Huang | 2019-04-30 | NTU experimental farm | B7C4R4 |
| 19TAI 84-B | Olram | Yung-Fen Huang | 2019-04-30 | NTU experimental farm | B7C4R6 |
| 19TAI 85-B | Shadow | Yung-Fen Huang | 2019-04-30 | NTU experimental farm | B7C4R7 |
| 19TAI 87-B | PI210075_11836 | Yung-Fen Huang | 2019-04-30 | NTU experimental farm | B8C1R5 |
| 20TW-19061 | PI182487 | Chung-Ying Ho | 2020-05-20 | NTU experimental farm | R12C14 |
| 20TW-SW1 | Swan | Chung-Ying Ho | 2020-06-01 | NTU experimental farm | R13C22 |
| 21TW-NTU1 | NTU Sel No. 1 | Chung-Ying Ho | 2021-02-21 | NTU experimental farm | R11C2 |
| 21TW-NTU3.2 | NTU Sel No. 1 | Chung-Ying Ho | 2021-02-21 | NTU experimental farm | R11C2 |
| 21TW-NTU4.2 | NTU Sel No. 1 | Chung-Ying Ho | 2021-02-21 | NTU experimental farm | R11C2 |

### Supplemental Table 2

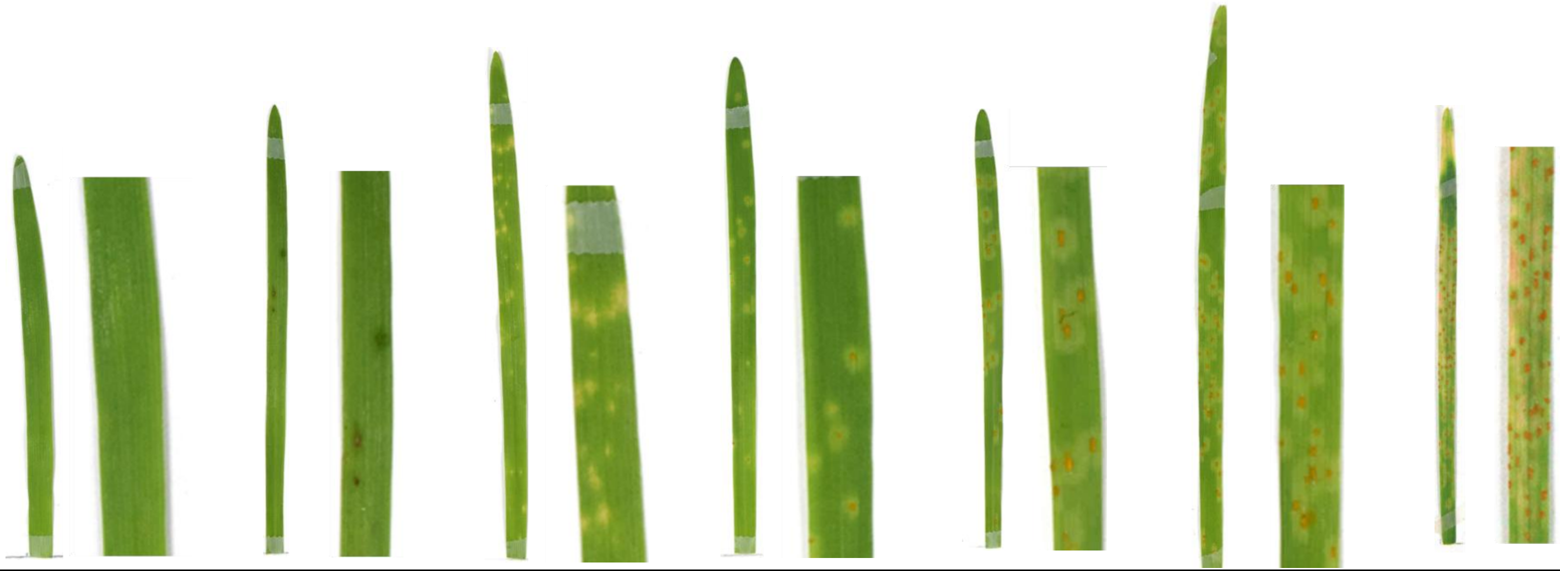

| Infection type | 0 | ; | ;c | 1 | 2 | 3 | 4 |
|---|---|---|---|---|---|---|---|
| Numeric scale | 0 | 1 | 2 | 3 | 4 | 5 | 6 |
| Infection response | Immune | Highly resistant | | Resistant | Moderately resistant | Moderately susceptible | Completely susceptible |
| Description | No uredinia or macroscopic symptom | No uredinia, but presence necrotic flecks | No uredinia; presence chlorosis, sometimes accompanied with necrosis | Few small uredinia surrounded by chlorosis, sometimes accompanied with necrosis | Small to medium sized uredinia in chlorotic areas, moderately distributed on the leaf, sometimes accompanied with necrosis | Medium to large sized uredinia in chlorotic areas | Large uredinia without necrosis or chlorosis. |

The scale was based on the rating scales developed by Murphy (1935), Chong et al. (2000), Nazareno et al. (2018), Miller et al. (2020), and our own observation. Photo scale: 1× for the left side and 2.5× for the right side.

### Supplemental Table 3

| Isolate | Species | Publication |
|---|---|---|
| TW-NTU01 | *Puccinia coronata* f. sp. *avenae* | Published here |
| 12NC29 | *Puccinia coronata* f. sp. *avenae* | Miller *et al*. 2018 |
| 12SD80 | *Puccinia coronata* f. sp. *avenae* | Miller *et al*. 2018 |
| *Pca*203 | *Puccinia coronata* f. sp. *avenae* | Henningsen *et al*. 2022 |
| SE-02 | *Puccinia graminis* f. sp. *avenae* | Lewis *et al*. 2018 |
| 98AG31 | *Melampsora larici-populina* | Pernaci *et al.* 2014 |
| *Pgt*21-0 | *Puccinia graminis* f. sp. *tritici* | Chen *et al*. 2017 |
| Ph560 | *Puccinia hordei* | Chen *et al*. 2019 |
| HN1907-1 | *Puccinia polysora* | Liang *et al*. 2022 |
| *Pst*130 | *Puccinia striiformis* f. sp. *tritici* | Cantu *et al*. 2011 |
| *Pt*76 | *Puccinia triticina* | Duan *et al*. 2022 |
